# Human Cytomegalovirus Immediate-Early 1 Protein Causes Loss of SOX2 from Neural Progenitor Cells by Trapping Unphosphorylated STAT3 in the Nucleus

**DOI:** 10.1101/308171

**Authors:** Cong-Cong Wu, Xuan Jiang, Xian-Zhang Wang, Xi-Juan Liu, Xiao-Jun Li, Bo Yang, Han-Qing Ye, Thomas Harwardt, Man Jiang, Hui-Min Xia, Wei Wang, William J. Britt, Christina Paulus, Michael Nevels, Min-Hua Luo

## Abstract

The mechanisms underlying neurodevelopmental damage caused by virus infections remain poorly defined. Congenital human cytomegalovirus (HCMV) infection is the leading cause of fetal brain development disorders. Previous work has linked HCMV to perturbations of neural cell fate, including premature differentiation of neural progenitor cells (NPCs). Here we show that HCMV infection of NPCs results in the loss of the SOX2 protein, a key pluripotency-associated transcription factor. SOX2 depletion maps to the HCMV major immediate-early (IE) transcription unit and is individually mediated by the IE1 and IE2 proteins. IE1 causes SOX2 down-regulation by promoting the nuclear accumulation and inhibiting the phosphorylation of STAT3, a transcriptional activator of SOX2 expression. Deranged signaling resulting in depletion of a critical stem cell protein is an unanticipated mechanism by which the viral major IE proteins may contribute to brain development disorders caused by congenital HCMV infection.

**IMPORTANCE:** Human cytomegalovirus (HCMV) infections are a leading cause of brain damage, hearing loss and other neurological disabilities in children. We report that the HCMV proteins known as IE1 and IE2 target expression of human SOX2, a central pluripotency-associated transcription factor that governs neural progenitor cell (NPC) fate and is required for normal brain development. Both during HCMV infection and when expressed alone, IE1 causes the loss of SOX2 from NPCs. IE1 mediates SOX2 depletion by targeting STAT3, a critical upstream regulator of SOX2 expression. Our findings reveal an unanticipated mechanism by which a common virus may cause damage to the developing nervous system and suggest novel targets for medical intervention.

## INTRODUCTION

Congenital human cytomegalovirus (HCMV) infection is the leading cause of birth defects worldwide. Approximately 1% of live newborns are infected *in utero* with this virus. At the time of birth, 5 to 10% of HCMV-infected newborn infants will exhibit signs of neurological damage, such as microcephaly, cerebral calcification and other abnormal findings (1–3). Among infected newborns that have no symptoms at birth, 10 to 15% subsequently develop central nervous system (CNS) disorders, including sensorineural hearing loss, mental retardation and learning disability (4–6). In addition, some authors have suggested that autism, language disorders and other more subtle changes in brain development might be related to congenital HCMV infection (7–9).

Although this virus can infect a wide range of organs *in vivo*, the fetal brain is regarded the principal target of HCMV infection that results in neurological manifestations (10–12). Due to exquisite host-specific tropism of this virus and the lack of animal models that faithfully recapitulate major characteristics of human infection, the pathogenesis of HCMV-associated disease in the developing CNS is largely unknown. However, recent advances in human neural progenitor cell (NPC) isolation and culture have provided an opportunity to study HCMV infection in a cell system relevant to fetal neuropathogenesis. Our previous studies and the work of others have shown that human NPCs are susceptible to HCMV infection and fully permissive to viral replication (13–19). HCMV infection of NPCs affects the cell fate by causing premature differentiation (14, 17, 18). Whole-genome analysis demonstrated that HCMV infection modulates the expression of NPC markers (14, 19) including sex-determining region Y (SRY)-box 2 (SOX2), a core transcriptional factor for stem cell self-renewal and pluripotency.

SOX2 is widely expressed in early neuroectoderm and neural progenitor cells during development (14, 19, 20) as well as in neural stem cells in the adult brain (20, 21). SOX2 missense or heterozygous loss-of-function mutations have been identified to cause ocular malformations, often manifesting as anophthalmia, microphthalmia, or coloboma. These symptoms may be accompanied by hearing loss, learning disability, or brain malformation (22–24). Familial recurrence of SOX2-associated anophthalmia has been observed (24). Moreover, the level of SOX2 expression plays an important role in sensory organ, including inner ear and retina, development. Groundbreaking research has demonstrated that forced SOX2 expression in fibroblasts, with or without additional factors, can generate induced pluripotent stem cells (25–27). In fact, ectopic SOX2 expression directs reprogramming of fibroblasts into neural stem or precursor cells (25, 28–30). SOX2 is also critical for maintenance of embryonic stem cells (ESCs). The SOX2 levels in ESCs are tightly regulated (31) and even small changes in expression can lead to differentiation (32, 33).

STAT3 is a member of the signal transducer and activator of transcription family (34) and is expressed in an activated form in the developing CNS as early as during initial NPC proliferation. This protein plays a dichotomous regulatory role in neuro- and gliogenesis (35, 36). STAT3 is activated through phosphorylation of tyrosine 705 (Y705) by receptor-associated kinases in response to various growth factors and cytokines including interleukin 6 (IL-6). Tyrosine phosphorylation leads to the nuclear accumulation of STAT3 homodimers, which act as DNA binding transcriptional activators of numerous target genes including SOX2. In ESCs that are differentiated into NPCs, STAT3 promotes cell fate commitment by activating the SOX2 promoter (37).

Here we investigate the effects of HCMV infection on SOX2 expression in human NPCs. We demonstrate that the HCMV 72-kDa immediate-early 1 (IE1) protein down-regulates SOX2 transcription and mediates depletion of the SOX2 protein from HCMV-infected NPCs. IE1 exerts its effect on SOX2 expression by inactivating the upstream regulator STAT3.

## RESULTS

### HCMV infection of NPCs causes down-regulation of SOX2 mRNA and protein

Previous work has established that human NPCs are fully permissive to HCMV infection (15, 38). As early as 4 h post infection (hpi), a significant decrease of SOX2 mRNA was observed in NPCs, and this decrease became more apparent as infection progressed (Fig 1A). In contrast, SOX2 protein levels did not change in these cells at early post-infection times (up to 12 hpi), but a gradual decrease was observed at late times (from 24 hpi) compared with mock-infected cells (Fig 1B). By 48 hpi, the SOX2 protein was clearly down-regulated, and by 96 hpi it was undetectable. The HCMV proteins IE1/IE2, UL44 and gB were also analyzed in this experiment to monitor IE, early, and late viral gene expression, respectively.

**Figure 1.**
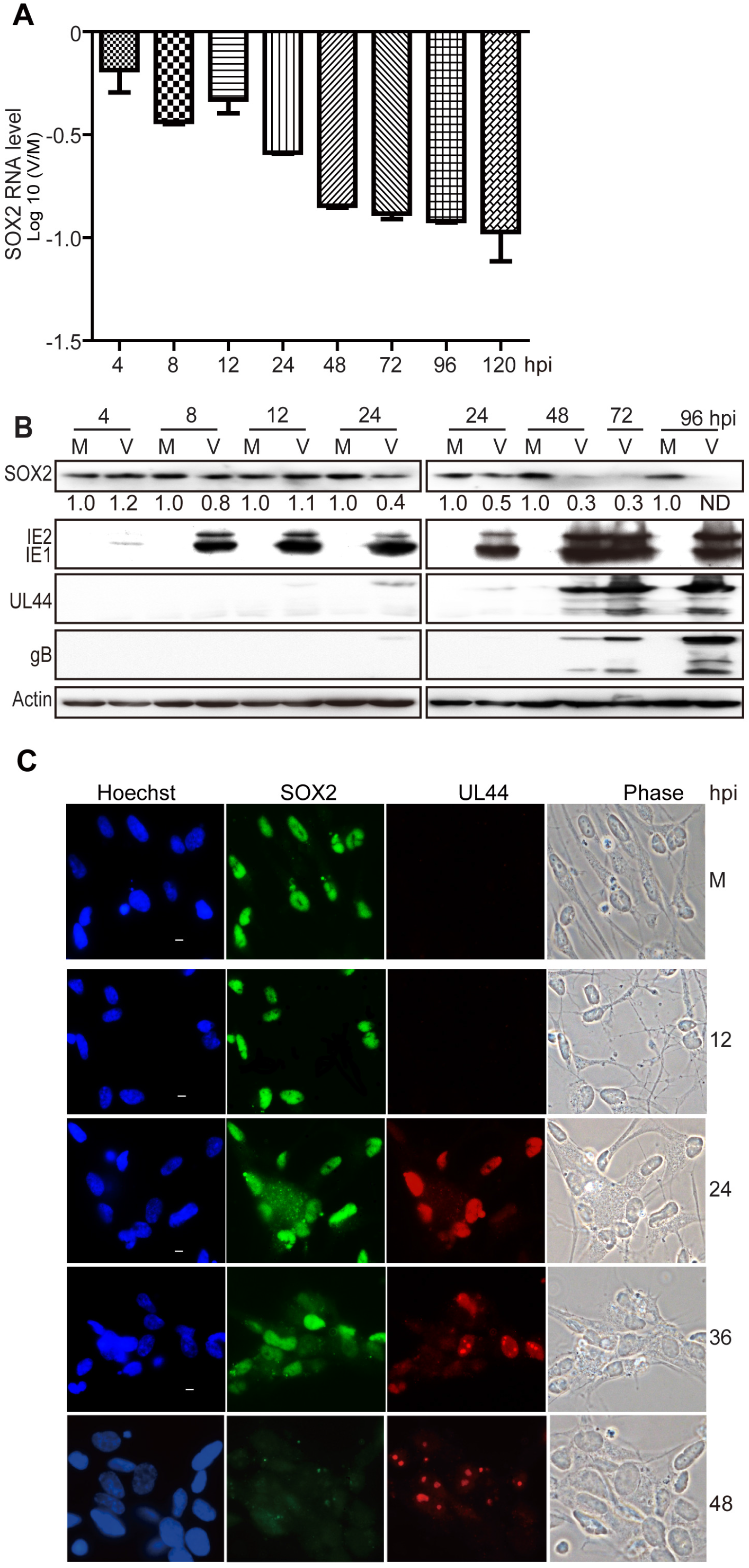
HCMV infection down-regulates SOX2 at the mRNA and protein level in NPCs. NPC monolayers were mock (M)- or virus (V)-infected with HCMV (TNWT) at an MOI of 3 and collected at the indicated times post infection for mRNA or protein analyses. (A) SOX2 mRNA levels during HCMV infection of NPCs. The levels of SOX2 mRNA normalized to GAPDH were determined by qRT-PCR at 4 to 120 hpi. Log_10_ values of virus-to-mock (V/M) ratios are given for each time point. Results shown are average ± standard deviation (SD) of data from three independent experiments, each conducted in triplicate. (B) SOX2 and viral protein levels during HCMV infection of NPCs. SOX2, IE1/IE2, UL44, and gB steady-state protein levels were determined by Western blotting at 4 to 96 hpi. Actin served as a loading control. The values listed below the blots indicate the relative SOX2 protein levels compared to corresponding mock controls following actin normalization. ND, not detectable. (C) Cellular distribution of SOX2 in relation to viral replication compartments during HCMV infection of NPCs. The distributions of SOX2 and UL44 were determined by indirect immunofluorescence assay at 12 to 48 hpi. NPCs grown on poly-D-lysine-coated coverslips were stained with antibodies against SOX2 (green) and UL44 (red), nuclei were counterstained with Hoechst 33342 (blue). Phase contrast images are also shown. Scale bars, 5μm.

In addition, the change in SOX2 protein levels was examined in relation to the formation and development of HCMV intranuclear replication compartments visualized via immunofluorescence staining of UL44 (Fig 1C). UL44 is the processivity factor of the HCMV DNA polymerase and is first expressed in the early phase of infection. Initially, UL44 was evenly distributed across the infected nucleus, then it formed multiple small foci (36 hpi) representing early viral replication compartments, and finally the small foci merged into bipolar foci (48 hpi) or single large late replication compartments (72 hpi, data not shown). In mock-infected NPCs, SOX2 exhibited its typical intense and diffuse nuclear staining. As expected, the SOX2 signal decreased during the course of HCMV infection inversely relating to UL44 foci development. In fact, SOX2 became dispersed and largely disappeared from late HCMV-infected NPCs.

These observations indicate the presence of a highly effective mechanism for SOX2 depletion in HCMV-infected NPCs.

### SOX2 depletion from HCMV-infected NPCs requires *de novo* viral protein synthesis

To determine if SOX2 down-regulation is dependent on HCMV gene products expressed *de novo* from the infecting viral genome, NPCs were exposed to infectious HCMV or UV-inactivated virus and SOX2 expression was analyzed. As expected, SOX2 expression was suppressed at both the RNA and protein level in infections with the untreated virus (Fig 2A and 2B). In contrast, UV-inactivated virus failed to suppress the expression of SOX2 mRNA (Fig 2A) and protein (Fig 2B). Similar levels of glycoprotein B (gB) derived from the input virus were detected at 12 hpi with both untreated and UV-treated HCMV, confirming that cells had been exposed to equal amounts of infecting virus (Fig 2B). However, *de novo* synthesis of viral proteins was undetectable in infections with UV-treated virus confirming successful inactivation. In another experiment, NPCs were pretreated with cycloheximide (CHX) and then infected with HCMV for 16 h in the presence of CHX. Down-regulation of SOX2 mRNA induced by HCMV infection was significantly diminished by CHX treatment (Fig 2C). No obvious change in SOX2 protein levels was observed by 16 hpi (data not shown), consistent with the results from Fig 1 and previous reports [14, 19].

**Figure 2.**
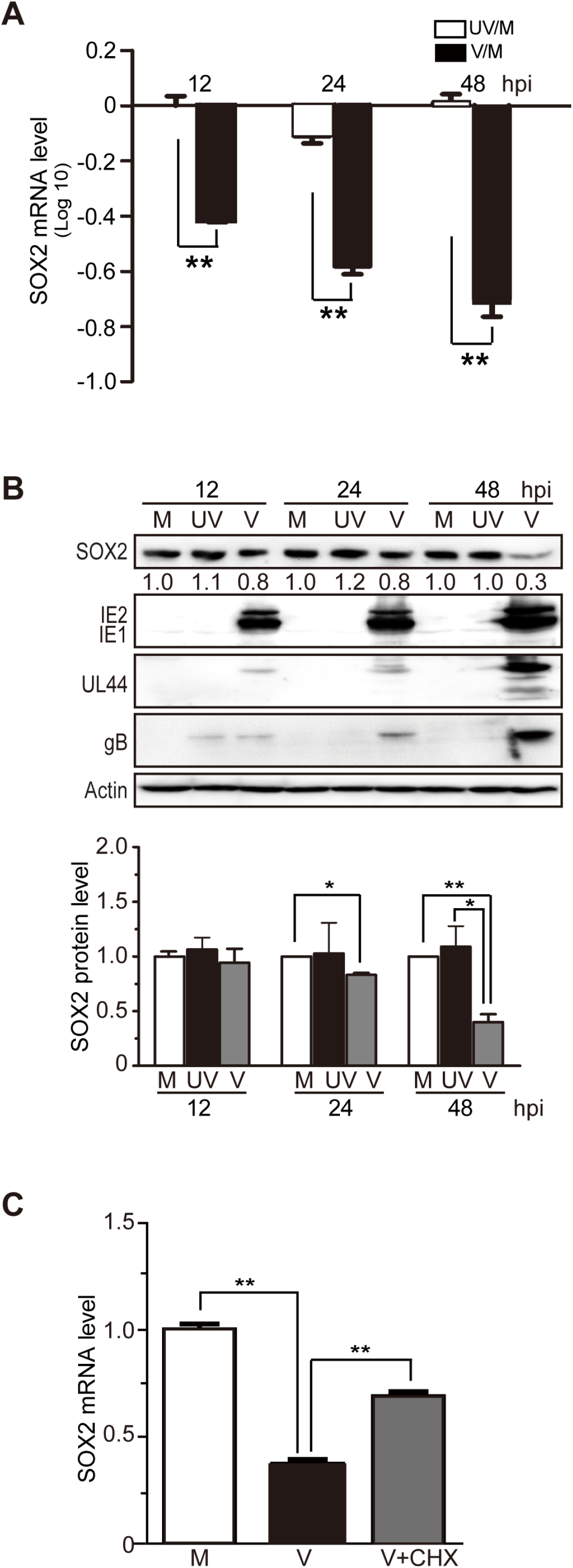
*De novo* synthesis of HCMV proteins is required for SOX2 down-regulation in NPCs. Comparison of HCMV infections with active virus (V) and UV-inactivated virus (UV). NPCs were mock-infected (M) or infected with active or UV-irradiated TNWT at an MOI of 3 and collected at the indicated times post infection. (A) SOX2 mRNA levels normalized to GAPDH were determined by qRT-PCR. Log_10_ values of inactivated virus-to-mock (UV/M) and active virus-to-mock (V/M) ratios are shown. Data from three independent experiments were analyzed by one way ANOVA, and results are presented as average ± SD. **, p ≤ 0.01. (B) Levels of SOX2 and representative viral proteins (IE1/IE2, UL44, and gB) were determined by Western blotting. Actin served as a loading control. The values listed below the blots indicate the relative SOX2 protein levels compared to corresponding mock controls following actin normalization. Data were from three independent experiments, results are presented as average ± SD. *, p ≤0.05; **, p ≤ 0.01. (C) Effect of protein synthesis inhibition by CHX treatment on SOX2 mRNA levels. NPCs were pretreated with CHX for 1 h prior to infection, and then mock (M)- or virus (V)-infected with HCMV (TNWT) at an MOI of 3. Cells were collected at 16 hpi for analysis of SOX2 mRNA by qRT-PCR. Data from three independent experiments were analyzed by one way ANOVA, and results are presented as average ± SD. **, p ≤ 0.01.

These results indicate that one or more viral proteins expressed *de novo* upon HCMV infection are required for SOX2 down-regulation in NPCs.

### HCMV major IE proteins mediate SOX2 down-regulation in NPCs

Transcription of SOX2 is suppressed as early as 4 hpi, a time when virion constituents and IE gene products are the only viral components present in the infected cell nucleus. To test whether IE or tegument proteins are sufficient for SOX2 down-regulation, the major IE proteins (IE1 and IE2) and the most abundant tegument protein (pp65) were individually expressed in NPCs using nucleofection. In agreement with the conclusion that newly synthesized viral proteins mediate SOX2 down-regulation (Fig 2), SOX2 levels did not change in NPCs transiently expressing pp65 compared to control cells (Fig 3A). Thus, pp65 was used as a negative control in subsequent assays. Expression of the entire major IE transcription unit, including the IE1 and IE2 proteins, resulted in significant reduction of SOX2 protein levels (Fig 3A). To discriminate between effects on SOX2 expression mediated by IE1 or IE2, each of the two proteins was individually expressed in NPCs. Following transfection of 2 or 5 μg IE1 expressing plasmid, the relative SOX2 mRNA levels were reduced to 61 or 23% of control levels, and the corresponding protein levels were reduced to 84 or 33%, respectively (Fig 3B). Transfection of 2 or 5 μg IE2 expressing plasmid reduced the relative SOX2 mRNA levels to 71 or 31% of control levels and the protein levels to 88 or 54%, respectively (Fig 3C).

**Figure 3.**
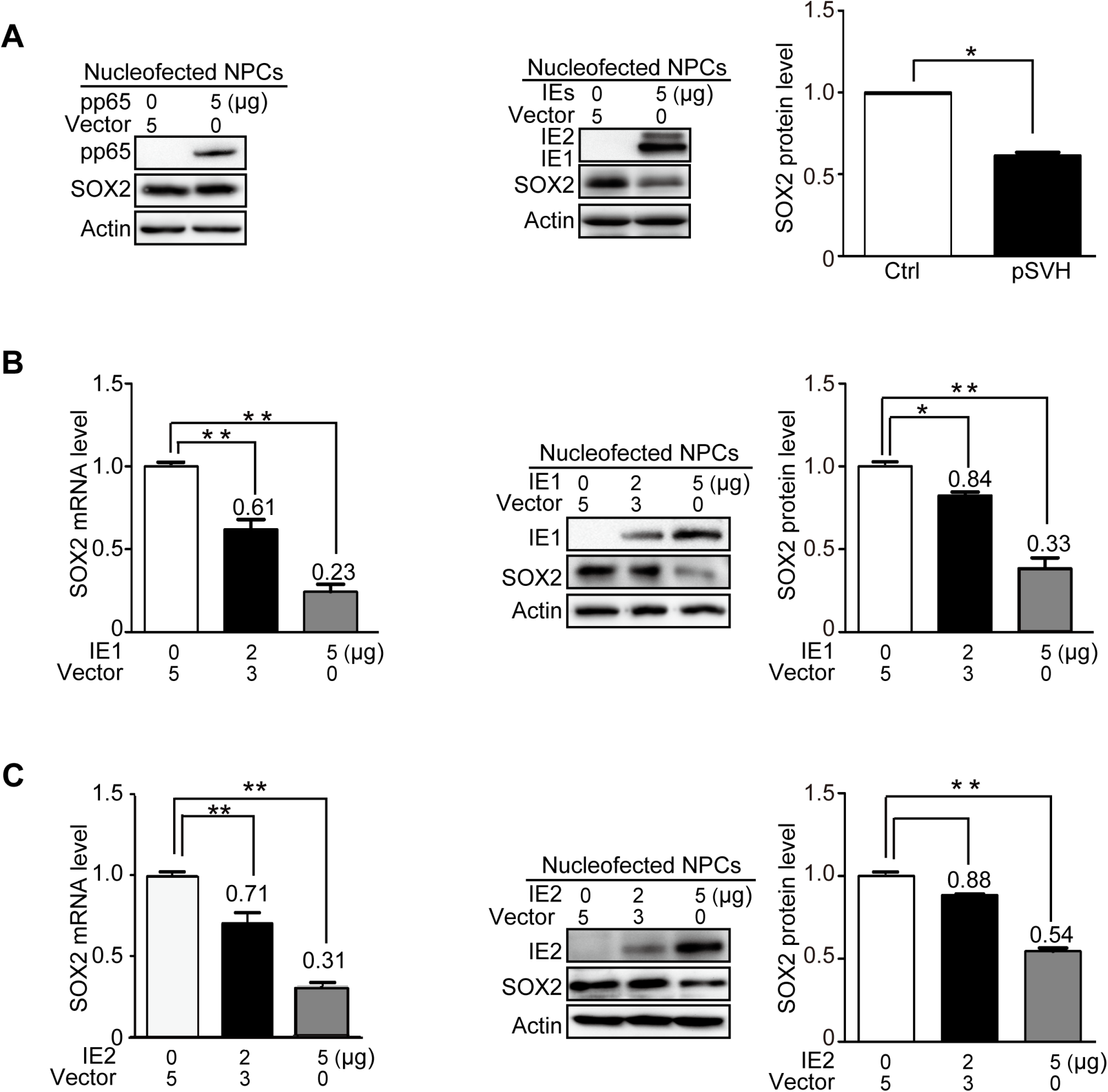
HCMV major IE proteins down-regulate SOX2 mRNA and protein in NPCs. HCMV pp65, IE1, IE2 or IE1 and IE2 combined were transiently expressed in NPCs following nucleofection. Samples were collected at 48 h post transfection for mRNA (qRT-PCR) or protein (Western blotting) analysis. Actin served as a loading control. The relative level of SOX2 protein compared to corresponding controls following actin normalization. Data from three independent experiments were analyzed by one way ANOVA, and results are presented as average ± SD. *, p ≤ 0.05; **, p ≤ 0.01. (A) Effect of pp65 and major IE proteins (IE1 and IE2) on SOX2 protein levels. Left: NPCs were transfected with 5 μg pcDNA3.0 (vector) or pcDNA3-pp65. Right: NPCs were transfected with 5 μg pcDNA3-pp65 as control (vector) or pSVH (IE1 and IE2). (B) Effect of IE1 on SOX2 mRNA and protein levels. NPCs were transfected with 5 μg pcDNA3-pp65 (vector), or 2 to 5 μg pcDNA3-IE1. (C) Effect of IE2 on SOX2 mRNA and protein levels. NPCs were transfected with 5 μg pcDNA3-pp65 (vector), or 2 to 5 μg pcDNA3-IE2.

These results indicate that the HCMV IE1 and IE2 proteins both play a significant role in down-regulating SOX2 expression. However, IE1 appeared to affect SOX2 more efficiently than IE2. Therefore, and because of the lethal phenotype of IE2-deficient viruses (39, 40), we focused our study on the role of IE1 in SOX2 regulation in present study.

### HCMV IE1 is required for depletion of SOX2 during infection of NPCs

HCMV infection and IE1 expression were each sufficient to efficiently down-regulate SOX2 at the mRNA and protein level in NPCs. To confirm the requirement of IE1 for SOX2 depletion during HCMV infection, the expression of IE1 was knocked down using shRNAs targeting IE1 exon 4 sequences. First, human embryonic lung (HEL) cells were transduced with lentiviruses expressing shRNAs directed against three different IE1-specific sequences, sh-1 to −3, or a scrambled shRNA control (scr). The transduced cells were subsequently synchronized and infected with HCMV. Analysis of IE1 protein levels determined at 24 and 48 hpi showed that sh-2 was the most efficient among the three tested shRNAs in reducing the levels of the IE1 protein (Fig 4A). To confirm the knock-down efficiency of sh-2, another experiment was performed where both IE1 mRNA and protein were analyzed at 24, 48 and 72 hpi. Again, sh-2 knocked down IE1 by >60% at the RNA level and 50% at the protein level through the course of infection (Fig 4B).

**Figure 4.**
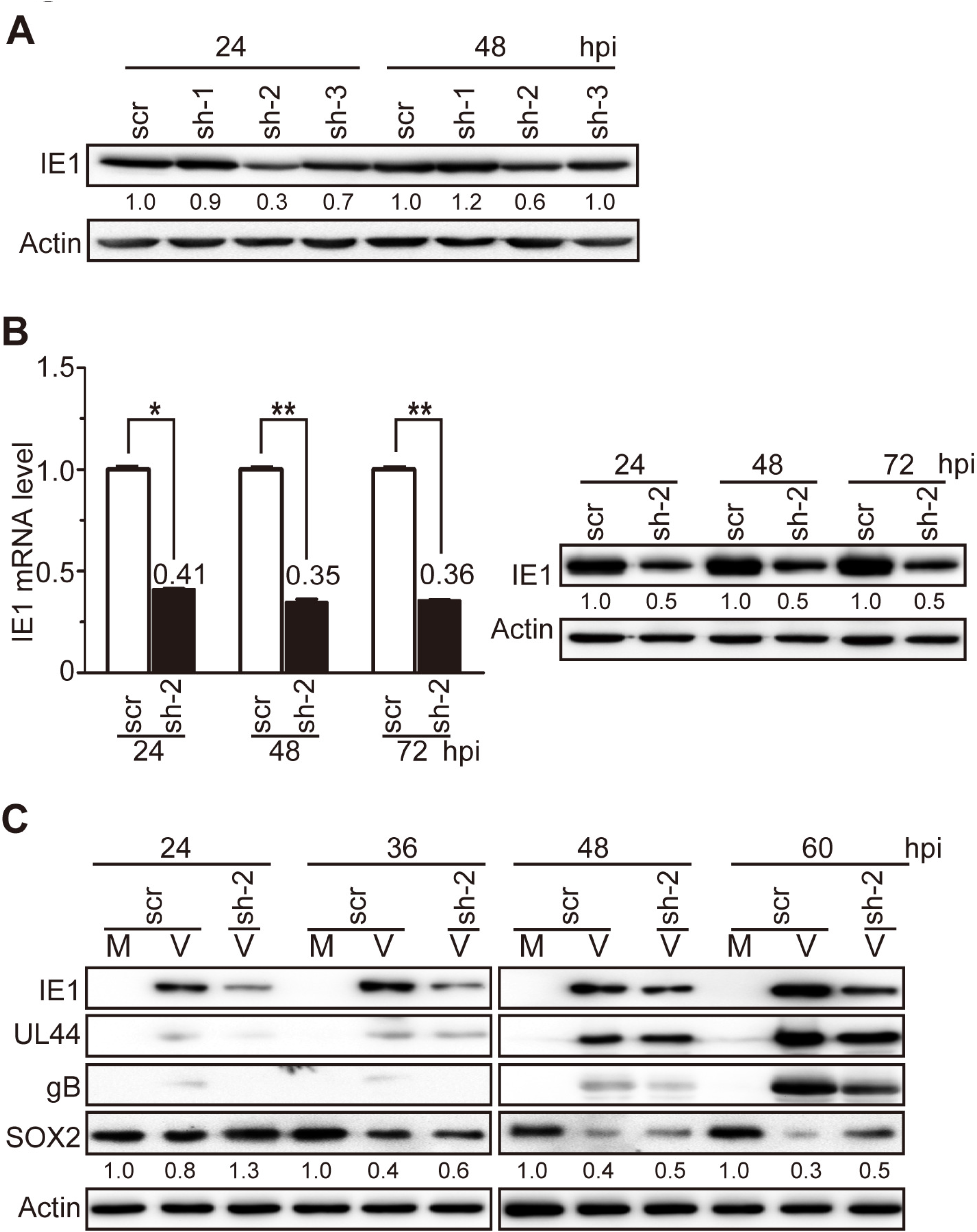
IE1 knock-down attenuates HCMV-induced SOX2 down-regulation in NPCs. (A) IE1-directed knock-down efficiency of candidate shRNAs in HEL cells. HEL cells were transduced with lentiviruses expressing shRNA-IE1-1 (sh-1), shRNA-IE1-2 (sh-2), shRNA-IE1-3 (sh-3), or shRNA-scramble (scr). At 48 h post transduction, cells were infected with HCMV (TNWT) at an MOI of 1 and collected at 24 or 48 hpi. IE1 protein levels were determined by Western blotting. Actin served as a loading control. (B) IE1-directed knockdown efficiency of sh-2 in HEL cells. HEL cells transduced with sh-2 or scr expressing lentiviruses were infected with HCMV (TNWT) as in (A). Left: IE1 mRNA levels were determined by qRT-PCR at the indicated times post infection; data from two independent experiments were analyzed by one way ANOVA, and results are presented as average ± SD. *, p ≤ 0.05; **, p ≤ 0.01. Right: protein levels were determined accordingly at the indicated times by Western blotting. (C) Effect of sh-2 on IE1 and SOX2 expression in NPCs. NPCs were transduced with sh-2 or scr expressing lentiviruses, cultured for 48 h, reseeded at a density of 3×10^6^ cells/dish, and mock (M)- or virus (V)-infected with TNWT at an MOI of 1. Protein levels of IE1, UL44, gB and SOX2 at the indicated times post infection are shown. Actin served as a loading control.

Following the successful IE1 knock-down in HEL cells, NPCs were transduced with lentiviruses expressing sh-2 or scr. The transduced NPCs were infected with HCMV and collected at 24 to 60 hpi. Since 24 hpi was previously shown to be the “turning point” of SOX2 protein levels (Fig 1), this time point was chosen as the earliest. As shown in Fig 4C, SOX2 protein levels gradually decreased as infection progressed, consistent with Fig 1. Significant knock-down of IE1 was observed at all tested time points in NPCs expressing sh-2 compared with scr expressing cells. SOX2 down-regulation was alleviated in HCMV-infected NPCs expressing sh-2 compared to cells expressing scr, most notably at 60 hpi.

Although the IE1 knock-down was incomplete and linked to reduced levels of other viral proteins (UL44 and gB), these results support the idea that IE1 contributes to SOX2 down-regulation during HCMV infection of NPCs.

### HCMV infection and IE1 expression inhibit STAT3 tyrosine phosphorylation and relocalize unphosphorylated STAT3 to the nuclei of NPCs

So far, our experiments have indicated that HCMV IE1 down-regulates SOX2, but a mechanism through which the viral protein alters expression of the cellular stem cell factor has not been identified. Results from co-immunoprecipitation and yeast two hybrid assays failed to provide evidence for a physical interaction between IE1 and SOX2 (data not shown). Since SOX2 expression was affected at both the protein and RNA level, it also seemed more likely that IE1 interferes with transcription rather than protein stability of SOX2. One key transcription factor regulating SOX2 expression is STAT3 (37), which has been recently identified as a physical interaction partner of IE1 (41, 42).

To investigate whether HCMV infection and IE1 expression affect SOX2 levels via modulation of STAT3 activation, the levels of total and tyrosine (Y705)-phosphorylated STAT3 (pSTAT3) were determined in infected NPCs. HCMV infection dramatically suppressed STAT3 tyrosine phosphorylation as early as 4 hpi, and pSTAT3 was maintained at low levels throughout the course of infection as compared to mock-infected NPCs. In contrast, the total STAT3 levels showed little if any changes during infection (Fig 5A).

**Figure 5.**
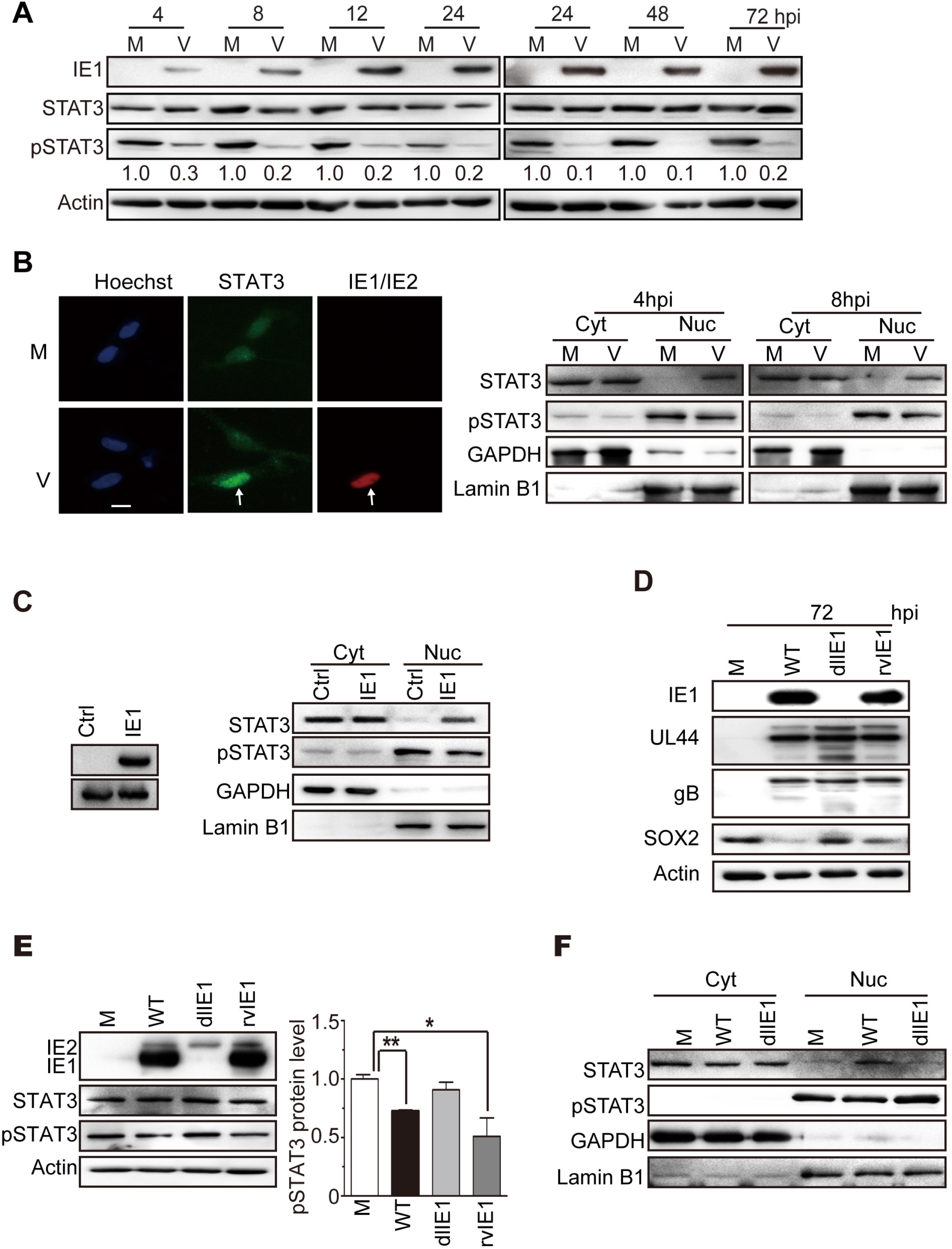
HCMV infection or IE1 expression inhibit STAT3 tyrosine phosphorylation and promote nuclear accumulation of unphosphorylated STAT3 in NPCs. (A) Inhibition of STAT3 tyrosine (Y705) phosphorylation by HCMV infection. NPCs were mock (M)- or virus (V)-infected with TNWT at an MOI of 3 and collected at the indicated times post infection. The protein levels of IE1, pSTAT3 and total STAT3 were determined by Western blotting. Actin served as a loading control. (B) Trapping STAT3 in nuclear by HCMV infection. NPCs were mock (M)- or virus (V)-infected with TNWT at an MOI of 1. Left: For indirect immunofluorescence analysis, NPCs on coverslips collected at 8 hpi were stained with antibodies against STAT3 (green) or IE1/IE2 (red), and nuclei were counterstained with Hoechst 33342 (blue). Infected (IE1/IE2 positive) cell is indicated with a white arrow. Scale bar, 10 μm. Right: For cellular fractionation analysis, fractions enriched in cytoplasmic (Cyt) or nuclear (Nuc) proteins were prepared from cells collected at 4 or 8 hpi. Protein levels of pSTAT3 and total STAT3 in each fraction were determined by Western blotting. GAPDH and lamin B1 served as controls for the Cyt and Nuc fraction, respectively. (C, D and E) Inhibition of tyrosine phosphorylation and nuclear sequestration of unphosphorylated STAT3 by IE1. Fractions enriched in cytosolic (Cyt) or nuclear (Nuc) proteins or total cell extracts were prepared. (C) For transient transfection analysis, NPCs were transfected with pcDNA3-IE1 or empty vector (Ctrl). Protein levels of IE1, pSTAT3, and total STAT3 were determined by Western blotting. GAPDH and lamin B1 served as controls for the Cyt and Nuc fraction, respectively. For HCMV infection analysis, NPCs were mock-infected (M) or infected with TNWT, TNdlIE1, or TNrvIE1 viruses at an MOI of 10. The levels of the indicated viral and cellular proteins in whole cell extracts (D, E) or in the Cyt and Nuc fractions (F) were determined by Western blotting. Actin, GAPDH and lamin B1 served as controls for total extracts or Cyt and Nuc fractions, respectively.

STAT3 is a nucleocytoplasmic shuttling protein which is efficiently exported from the nucleus in its unphosphorylated form (43). Unphosphorylated STAT3 is therefore typically located in the cytoplasm or distributed across both the nuclear and cytoplasmic compartments. However, upon tyrosine phosphorylation pSTAT3 efficiently accumulates in the nucleus (37, 44, 45). To define the subcellular distribution of STAT3 in HCMV-infected NPCs, we performed immunofluorescence analysis. NPCs on coverslips were infected with HCMV (MOI=1), collected at 8 hpi, and stained for STAT3 and IE1/IE2. STAT3 was mainly confined to the nuclei of virus-infected cells, while in mock-infected cells the STAT3 signal was more diffuse and cytoplasmic as well as nuclear (Fig 5B). This observation was consistent with our results from HCMV-infected fibroblasts (41, 42). To further investigate STAT3 subcellular localization in HCMV-infected NPCs, the levels of pSTAT3 and total STAT3 in cytoplasmic and nuclear compartments separated by cellular fractionation were analyzed at 4 and 8 hpi. GAPDH and lamin B1 served as loading controls for the cytoplasmic and nuclear compartments, respectively, confirming successful fractionation. As expected, pSTAT3 was present predominantly in the nucleus of both mock- and HCMV-infected cells. Compared to mock-infected NPCs, pSTAT3 levels were reduced in both the cytoplasm and nucleus of HCMV-infected cells at 4 and 8 hpi. Total STAT3 resided mainly in the cytoplasm, but the nuclear signal was much stronger in HCMV-infected compared with mock-infected cells at both tested time points (Fig 5B). These results indicate that HCMV infection reduces the levels of activated STAT3 (pSTAT3) and promotes the nuclear accumulation of unphosphorylated STAT3 at very early times of infection in NPCs.

IE proteins are thought to be the only HCMV gene products expressed at 4 to 8 hpi, when STAT3 is relocalized to the nucleus and its phosphorylation is inhibited. To test whether IE1 was responsible for the observed effects on STAT3, the viral protein was transiently expressed in NPCs. IE1 expression was sufficient to down-regulate pSTAT3 levels and to sequester an unphosphorylated form of the cellular protein in the nuclear compartment (Fig 5C). To further confirm the role of IE1 in altering STAT3 phosphorylation and intracellular localization, NPCs were mock-infected or infected with WT, dlIE1, or rvIE1 viruses. The levels of viral proteins (UL44, gB) were similar, except that IE1 was absent and down-regulation of SOX2 was diminished in the infection with dlIE1 (Fig 5D). Compared with mock-infected cells, the pSTAT3 levels were reduced in WT- and rvIE1-, but not in dlIE1-infected cells. Again, the overall steady-state STAT3 levels were rather constant across all mock- and HCMV-infected samples (Fig 5E). Finally, based on cellular fractionation analysis, pSTAT3 was located in the nuclei of both mock- and WT- or dlIE1-infected cells. Total STAT3 localized mainly in the cytoplasm of mock- and dlIE1-infected cells, while increased nuclear localization was observed in WT-infected NPCs (Fig 5F).

These results demonstrate that IE1 is both sufficient and required for the inhibition of tyrosine phosphorylation and nuclear sequestration of unphosphorylated STAT3 observed during HCMV infection of NPCs.

### SOX2 down-regulation in HCMV-infected NPCs results from IE1-mediated inhibition of STAT3 activation

The results thus far obtained are all consistent with a mechanism in which IE1 mediates SOX2 down-regulation by limiting STAT3 activation. To further test this possibility, we performed several different experiments. We first treated NPCs with CTS, a chemical inhibitor of STAT3 tyrosine phosphorylation (46). In the presence of CTS, pSTAT3 levels (but not steady-state STAT3 levels) markedly decreased as a function of time of treatment coinciding with gradual loss of SOX2 (Fig 6A). Then, STAT3 expression was silenced using RNA interference. Two different shRNAs, shSTAT3-1 and shSTAT3-2, knocked down STAT3 expression with different efficiencies compared to non-specific shRNAs (shLuci and shDsRed). The extent of STAT3 silencing correlated with differential reduction in pSTAT3 levels, in turn correlating with the levels of SOX2 (Fig 6B).

**Figure 6.**
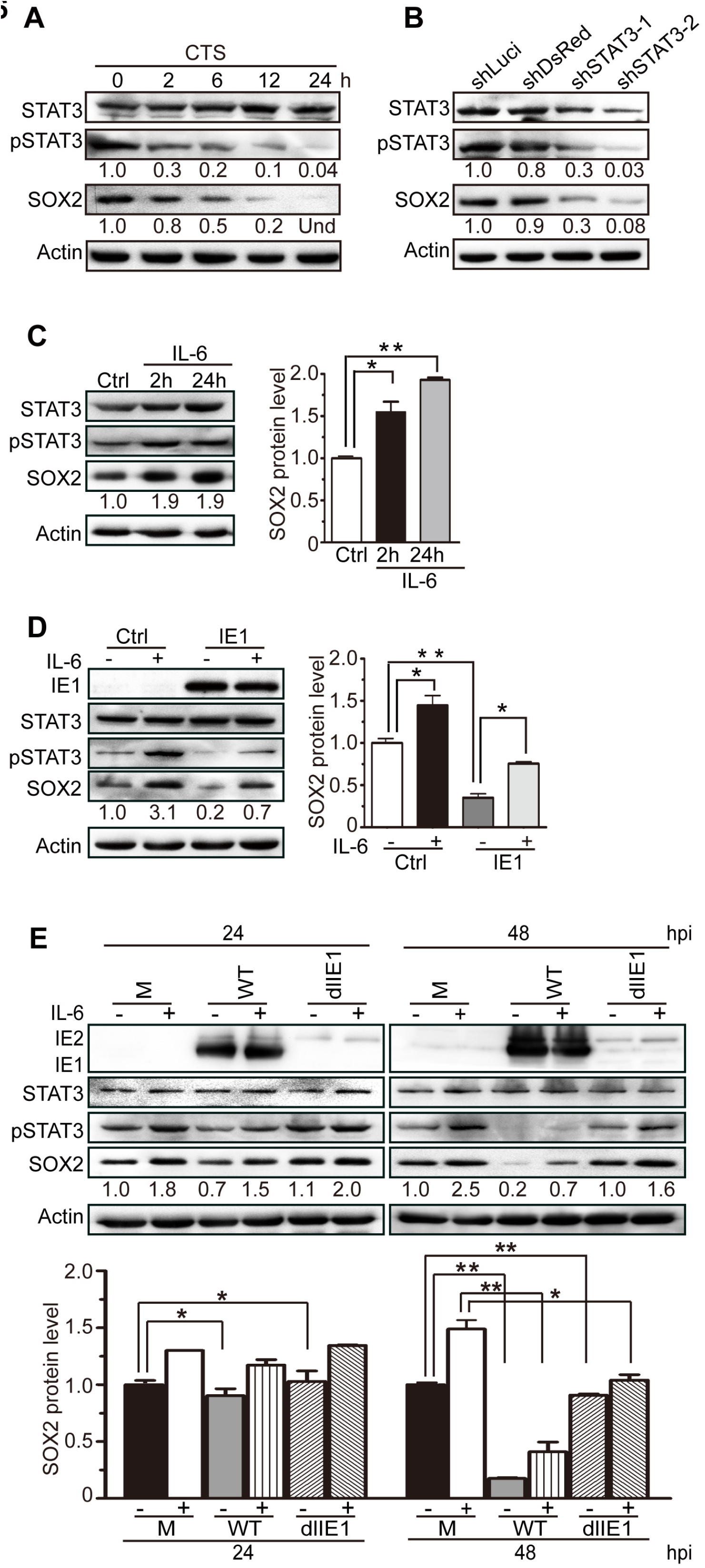
SOX2 expression strictly depends on pSTAT3, and IE1 mediates SOX2 depletion by inhibiting STAT3 activation. (A) Inhibition of STAT3 correlates with suppression of SOX2 expression. NPCs were treated with the chemical inhibitor cryptotanshinone (CTS) for the indicated times. The protein levels of total STAT3, pSTAT3, and SOX2 were determined by Western blotting. Actin served as a loading control. Und, undetectable. (B) Silencing of STAT3 correlates with suppression of SOX2 expression. NPCs were transduced with lentiviruses Tet-pLKO-puro-shLuci (shLuci), Tet-pLKO-puro-shDsRed (shDsRed), Tet-pLKO-puro-shSTAT3-1 (shSTAT3-1) or Tet-pLKO-puro-shSTAT3-2 (shSTAT3-2) and treated with doxycycline for 48 h to induce shRNA expression. The protein levels of total STAT3, pSTAT3, and SOX2 were determined by Western blotting. Actin served as a loading control. (C) IL-6-mediated activation STAT3 correlates with induction of SOX2 expression. NPCs were treated with IL-6 for 2 or 24 h. The protein levels of pSTAT3, total STAT3 and SOX2 were monitored by Western blotting. Actin served as a loading control. Data from three independent experiments were analyzed by one way ANOVA, and results are presented as average ± SD. *, p ≤ 0.05; **, p ≤ 0.01. (D) IL-6 counteracts IE1-dependent SOX2 down-regulation in transiently transfected NPCs. NPCs were transfected with pcDNA3-IE1, treated with IL-6 for 4 h or left untreated and collected at 48 h post transfection. The protein levels of IE1, total STAT3, pSTAT3, and SOX2 were determined by Western blotting. Actin served as a loading control. Data from three independent experiments were analyzed by one way ANOVA, and results are presented as average ± SD. *, p ≤ 0.05; **, p ≤ 0.01. (E) IL-6 counteracts IE1-dependent SOX2 down-regulation in HCMV-infected NPCs. NPCs were infected with TNWT or TNdlIE1 viruses at an MOI of 10, treated with IL- 6 for 4 h or left untreated, and collected at 48 hpi. The protein levels of IE1/IE2, total STAT3, pSTAT3, and SOX2 were determined by Western blotting. Actin served as a loading control. Data from three independent experiments were analyzed by one way ANOVA, and results are presented as average ± SD. *, p ≤ 0.05; **, p ≤ 0.01

In addition to the experiments involving inhibition of STAT3, we set out to study the IE1-STAT3-SOX2 axis by activating STAT3 signaling. To this end, we first treated NPCs with IL-6, a major agonist of STAT3 signaling (47). IL-6 treatment led to a marked increase in pSTAT3 without significantly affecting the overall STAT3 protein levels. Concurrent with increased STAT3 tyrosine phosphorylation, there was also a rise in SOX2 levels. These data were consistent with the finding that pSTAT3 promotes SOX2 expression in NPCs (Fig 6C). Next, we transiently expressed IE1 in NPCs, treated the cells with IL-6 for 4 h or left them untreated, and analyzed the levels of IE1, pSTAT3, total STAT3 and SOX2. Both pSTAT3 and SOX2 were strongly up-regulated by IL-6, but down-regulated by IE1, and IL-6 efficiently counteracted IE1-mediated down-regulation of pSTAT3 and SOX2. Again, the total steady-state STAT3 levels did not significantly change upon IL-6 treatment or IE1 expression (Fig 6D). Finally, to further investigate the link between IE1 expression, STAT3 activation and SOX2 regulation in the context of HCMV infection, NPCs were mock-infected or infected with HCMV WT or dlIE1. The infected NPCs were treated with IL-6 for 4 h or left untreated prior to collection at 24 and 48 hpi. As expected, IL-6 treatment led to elevated levels of pSTAT3 and SOX2 in mock-, WT- and dlIE1-infected NPCs. HCMV WT infection reduced the pSTAT3 levels slightly at 24 hpi and dramatically at 48 hpi. In contrast, no obvious decrease of pSTAT3 was observed following dlIE1 infection. Concordantly, SOX2 protein levels fell substantially in WT infection at 48 hpi, but remained constant in dlIE1 infection at the same time point. Again, IL-6 treatment counteracted IE1-dependent down-regulation of both pSTAT3 and SOX2 (Fig 6E).

In summary, these data demonstrate that SOX2 expression in NPCs strictly depends on pSTAT3, and that there is a causal link between SOX2 down-regulation by HCMV and inhibition of STAT3 activation by IE1, including inhibition of tyrosine phosphorylation and trapping unphosphorylated STAT3 in nuclear.

## DISCUSSION

Proliferation, differentiation and migration of NPCs as well as synapse formation among mature neurons are all critical factors for fetal brain development and function (48–50). HCMV infection has been shown to induce neural cell loss and abnormal differentiation of NPCs (14, 15, 19). This was not only demonstrated *in vitro*, but also confirmed in a mouse model of congenital infection where the virus appeared to affect NPCs in the subventricular zone (12). Although HCMV is the leading cause of neurological damage in children, the mechanisms by which the virus perturbs NPC proliferation and differentiation have remained unclear.

SOX2 is a master controller of stem cells, strikingly illustrated by the fact that its overexpression can reprogram terminally differentiated fibroblasts to induced pluripotent stem cells (25–30). More specifically, SOX2 is a transcription factor essential for maintaining selfrenewal and pluripotency of ESCs and NPCs. SOX2 mutations cause symptoms resembling damage resulting from congenital HCMV infection, including neural development disorders accompanied by ocular malformation (22–24). In this study, we demonstrate that HCMV significantly down-regulates SOX2 mRNA levels in human primary NPCs from as early as 4 hpi, extending our previous observations (14, 38). The reduced mRNA levels subsequently lead to almost full depletion of the SOX2 protein from HCMV-infected cells at later times (24 to 96 hpi). The temporal delay between mRNA and protein down-regulation suggests that the SOX2 protein may be very stable in NPCs, in contrast to what has been reported for ESCs (51). The timing of changes in SOX2 mRNA levels and the observation that the down-regulation depends on *de novo* viral protein synthesis, pointed us to the HCMV major IE proteins as potential regulators of this stem cell factor. The 72-kDa IE1 and the 86-kDa IE2 protein subsequently proved to be sufficient to decrease SOX2 mRNA and protein levels in the absence of other viral proteins, together and individually. IE1 and IE2 are nuclear localized key regulators of viral and cellular transcription during infection (52–54). They have been extensively studied in fibroblasts, but there is little information on how these proteins behave in cell types more relevant to HCMV pathogenesis.

Our results demonstrate that the HCMV IE1 protein is not only sufficient, but also necessary for the reduction of SOX2 levels through the early-late stages of infection. As noted above, the down-regulation of SOX2 RNA and protein was also observed with ectopically expressed HCMV IE2, but this protein appeared to be less efficient than IE1 in this respect. We propose that IE2 also contributes to SOX2 down-regulation in the context of HCMV infection, although this remains to be formally shown. Due to the difficulties in working with IE2-deficient viruses (IE2 is essential for HCMV replication and difficult to complement), we focused on IE1 in this study. Our findings may come as a surprise in the light of two previous reports which concluded that HCMV infection or IE1 expression leads to increased instead of decreased SOX2 levels in human glioma stem-like cells, human glioblastoma cells and mouse glioma tissue (55, 56). The seemingly disparate findings might be due to differences in cell types or virus strains. However, in our hands, IE1 induced robust inhibition of STAT3 activation in all cell types tested so far including glioblastoma-derived cell lines (data not shown). Notably, the previous studies did not discriminate between the levels of SOX2 in IE1 expressing cells and IE1-negative cells in the same culture or tumor. Even if the STAT3 pathway is inhibited in HCMV-infected cells or cells ectopically expressing IE1, the secretome associated with these cells may trigger STAT3 signaling in IE1-negative bystander cells. In population analyses, this bystander effect may be detected as an overall increase in STAT3-dependent gene expression (42).

Our work also determines the molecular events underlying IE1-dependent SOX2 down-regulation, which appear to be intimately linked to JAK-STAT signaling. JAK-STAT (STAT1 and STAT3) signaling pathways play an important role in neural development by regulating NPC neurogenesis and gliogenesis (35, 36). IE1 markedly affects the activation state and subcellular localization of STAT3, a key regulatory protein known to affect the pluripotency of ESCs and initiate the commitment to NPC fate via transcriptional activation of SOX2 (37). IE1 inhibits tyrosine (Y705) phosphorylation and promotes nuclear accumulation of STAT3 without altering the protein’s overall steady-state levels. These observations closely recapitulate previous results in fibroblasts and are likely the consequence of a direct physical interaction between IE1 and STAT3 (41, 42). Although paradoxical on the face of it, decreased tyrosine phosphorylation and increased nuclear localization of STAT3 may be coupled. STAT proteins, including STAT3, continually shuttle between the cytoplasm and the nucleus. In fact, STAT3 is imported to the nucleus independent of tyrosine phosphorylation, and this is normally followed by export to the cytoplasm. Tyrosine phosphorylation transiently increases STAT3 nuclear accumulation due to sequestration by DNA binding or heterodimerization with other phosphorylated STAT proteins (43). Likewise, IE1 may bind to STAT3 passing through the nucleus. Nuclear sequestration may reduce the amounts of STAT3 undergoing export to the cytoplasm and, consequently, the pools of cytoplasmic STAT3 available for tyrosine phosphorylation by the corresponding (cytoplasmic) JAK family kinases. The expected consequences of depleting activatable STAT3 by nuclear sequestration are in line with the reduced pSTAT3 levels, restricted responsiveness to IL-6 and diminished expression of SOX2 we observed in HCMV-infected NPCs.

In summary, this study initially links the interaction between IE1 and STAT3 to SOX2 depletion and thereby identifies a novel pathway predicted to contribute to developmental neuropathogenesis caused by congenital HCMV infection.

## MATERIALS AND METHODS

### Ethics statement

Postmortem fetal brain tissues from different gestational age cases were obtained according to the approval notice from the Institutional Review Board (WIVH10201202) and the Guidelines for Biomedical Research Involving Human Subjects at Wuhan Institute of Virology, Chinese Academy of Sciences. The original source of the anonymized tissues was Zhongnan Hospital of Wuhan University (China). The cell isolation procedures and research plans were approved by the Institutional Review Board (IRB) (WIVH10201202) according to the Guidelines for Biomedical Research Involving Human Subjects at Wuhan Institute of Virology, Chinese Academy of Sciences. The need for written or oral consents was waived by IRB (57).

### Cells and cell culture

All cells were maintained at 37°C in a humidified atmosphere containing 5% CO_2_. NPCs were isolated as described previously (38, 58) and cultured in a 1:1 mix of growth medium and conditioned medium (14, 15, 19). The NPC growth medium was Dulbecco’s Modified Eagle Medium (DMEM)-F12 (Thermo Fisher Scientific) supplemented with 2 mM GlutaMAX (Thermo Fisher Scientific), 100 U/ml penicillin and 100 μg/ml streptomycin (Thermo Fisher Scientific), 50 μg/ml gentamycin (Sigma), 1.5 μg/ml amphotericin B (Thermo Fisher Scientific), 10% BIT 9500 (Stem Cell Technologies), 20 ng/ml epidermal growth factor (EGF, Prospec) and 20 ng/ml basic fibroblast growth factor (FGF, Prospec). Conditioned medium was collected from cultured NPCs and stored at −20°C after cell debris had been removed by centrifugation. NPCs were maintained as monolayers in fibronectin-coated dishes and seeded at a defined density into dishes coated with poly-D-lysine (50 μg/ml, Millipore) prior to infection. In order to induce STAT3 activation, NPCs were treated with carrier-free recombinant human IL-6 (183 ng/ml, Biolegend) for the indicated times before being collected for protein analysis. To inhibit STAT3 activation, NPCs were treated for the indicated times with 10 μM cryptotanshinone (CTS, Sigma), a small molecular inhibitor of STAT3 tyrosine (Y705) phosphorylation.

Human embryonic lung fibroblasts (HEL cells, maintained in the laboratory) and HEL cells immortalized by transfection with pCI-neo-hTERT (HELf cells, kindly provided by Dr. Chen, Columbia University), were cultured in Minimal Essential Medium (MEM, Thermo Fisher Scientific). Human embryonic kidney (HEK) 293T cells (CRL-11268) were grown in DMEM. Both MEM and DMEM were supplemented with 10% fetal bovine serum (FBS, Thermo Fisher Scientific), penicillin and streptomycin, and 2 mM L-glutamine (Thermo Fisher Scientific). For the establishment of an IE1 expressing HELf cell line, a lentivirus stock was prepared from plasmid pCDH-puro-IE1. A volume of 2 ml from this stock was used to transduce 1×10^6^ HELf cells. Transduced cells were cultured in normal medium for 2 days to allow for transgene expression, and then switched to selection medium containing 8 μg/ml puromycin (Sigma) for 2 weeks with medium changes every other day. Expression of IE1 was confirmed by Western blotting. The resulting HELf cell line stably expressing IE1 was designated HELf-IE1 and maintained in medium containing puromycin (4 μg/ml).

### Plasmids and nucleofection

Plasmids pcDNA3-IE1 and pcDNA3-IE2 were constructed by subcloning a *Bam*HI-*Bam*HI fragment containing the full-length coding sequence of the HCMV 72-kDa IE1 or 86-kDa IE2 protein, respectively, from pSG-IE1 or pSG-IE2 (kindly provided by Dr. Fortunato, University of Idaho), respectively, into the backbone of pcDNA3.0. Plasmid pcDNA3-pp65 was generated by cloning a fragment PCR-amplified from HCMV (Towne) cDNA between the *Bam*HI and *Eco*RI sites of pcDNA3.0 (57). Plasmid pSVH (kindly provided by Dr. Tang, Howard University) containing the entire HCMV major IE transcription unit (including IE1, IE2 and the major IE promoter-enhancer) was also used.

Plasmids expressing short hairpin RNAs (shRNAs) directed against HCMV (Towne) IE1 sequences were constructed based on lentiviral vector pLKO.1 puro (Addgene plasmid #8453). Three shRNAs targeting IE1 exon 4 and a scrambled shRNA not targeting any viral or human gene were designed (http://jura.wi.mit.edu/bioc/siRNAext), synthesized, and inserted between the *Age*I and *Eco*RI sites of pLKO.1 puro to generate pLKO.1-shRNA-IE1-1 (sh-1), pLKO.1 -shRNA-IE1 −2 (sh-2), pLKO.1-shRNA-IE1-3 (sh-3), and pLKO.1-scramble (scr), respectively. Sequences are listed in Table 1. For the establishment of HELf-IE1 cells (described above), a fragment containing the IE1 coding sequence was recovered from pSG-IE1 and cloned into the *Bam*HI site of pCDH-CMV-MCS-EF1-puro (System Biosciences). The resulting construct was designated pCDH-puro-IE1.

**Table 1.**
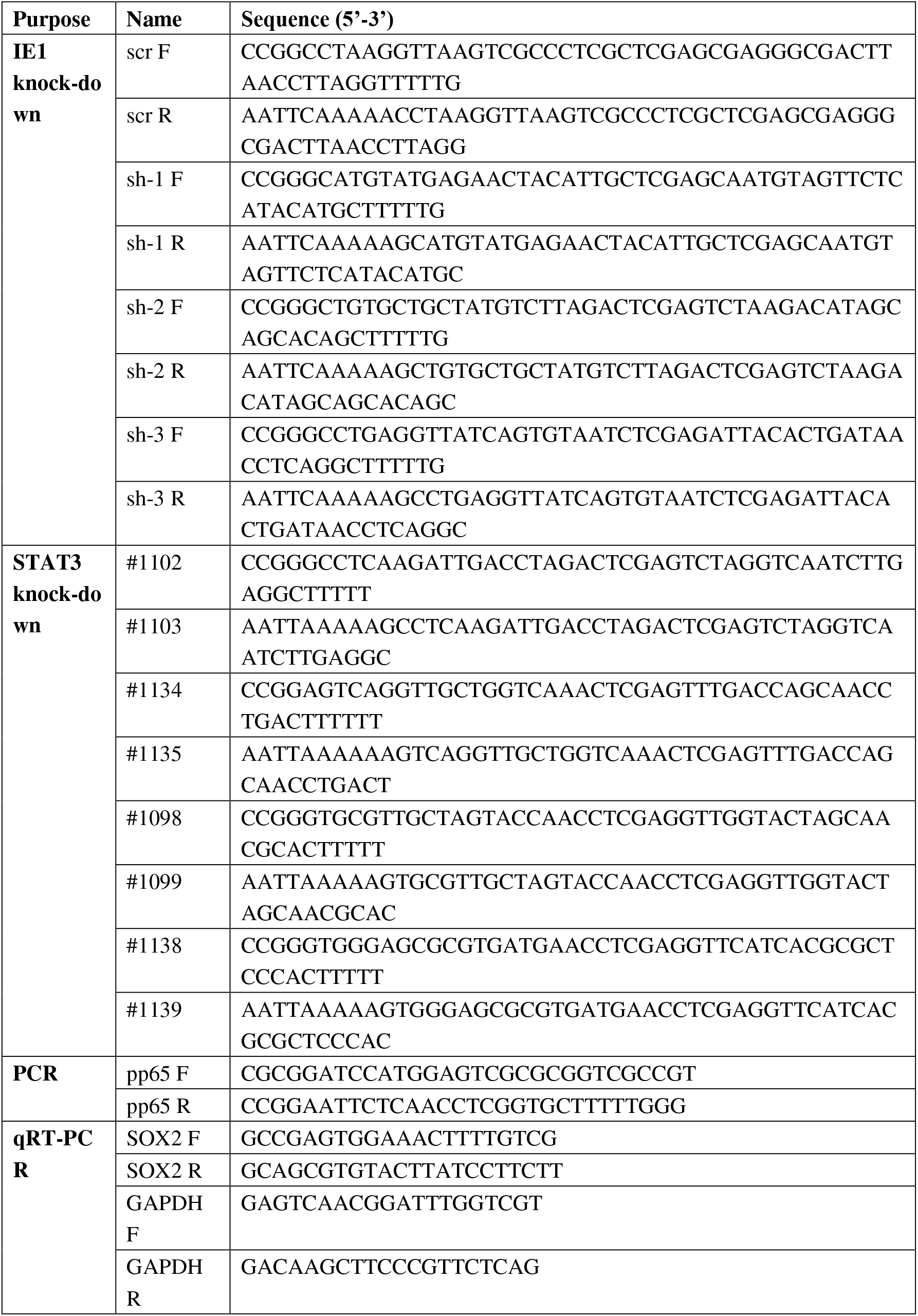
Oligonucleotides used in this study.

Plasmids expressing shRNAs directed against human STAT3 sequences were constructed based on lentiviral vector Tet-pLKO-puro (Addgene plasmid #21915). Two shRNA plasmids targeting STAT3, Tet-pLKO-puro-shSTAT3-1 and Tet-pLKO-puro-shSTAT3-2, were generated by inserting annealed oligonucleotides #1102 and #1103 or #1134 and #1135, respectively, between the *Eco*RI and AgeI sites of Tet-pLKO-puro. Likewise, plasmids Tet-pLKO-puro-shLuci and Tet-pLKO-puro-shDsRed expressing shRNAs directed against *Photinus* luciferase or *Discosoma* red fluorescent protein sequences, respectively, not present in human cells were constructed by inserting annealed oligonucleotides #1098 and #1099 or #1038 and #1039, respectively, between the *Eco*RI and *Age*I sites of Tet-pLKO-puro. Oligonucleotide sequences are listed in Table 1.

To transiently overexpress individual viral proteins, NPCs were transfected with plasmids pSVH, pcDNA3-IE1, pcDNA3-IE2, pmaxGFP (provided by Lonza to assess transfection efficiency), pcDNA3-pp65, or empty vector (pcDNA3.0) using Nucleofector technology (Lonza) according to the manufacturer’s instructions. In brief, 5×10^6^ NPCs were mixed with 100 μl Mesenchymal Stem Cell (MSC) Nucleofector Solution (82 μl Nucleofector Solution with 18 μl Supplement 1) and 5 μg pSVH, pcDNA3-IE1, pcDNA3-IE2, pmaxGFP, pcDNA3-pp65, or pcDNA3.0. The cell-DNA suspension was transferred to certified cuvettes, which were inserted into a Nucleofector II, and program A-033 was applied. Immediately following nucleofection, 500 μl NPC growth medium was added to the cuvette, and the sample was gently transferred to poly-D-lysine-coated dishes. After 24 h, the culture medium was replaced, and cells were analyzed at 48 h post transfection.

### HCMV preparation and infection

Enhanced green fluorescent protein (EGFP) expressing bacterial artificial chromosome-derived wild-type (T-BAC, herein referred to as TNWT), IE1-deficient (TNdlIE1), and revertant (TNrvIE1) variants of the HCMV Towne strain (ATCC-VR977) and the parental virus HCMV Towne strain (ATCC-VR977) were used in this study. The construction of TNdlIE1 and TNrvIE1 was described previously (59). The TNWT and TNrvIE1 viruses were propagated in HEL cells and titrated by plaque assay as described previously (60, 61). The TNdlIE1 mutant was propagated and titrated in HELf-IE1 cells. HCMV particles from infected cell supernatants were concentrated by ultracentrifugation, after removing the cell debris by high speed centrifugation, and resuspended in NPC growth medium to avoid any potential undesired effects induced by components (including FBS) in the medium used for virus preparation (61). UV-inactivated HCMV was prepared by exposure to 6000 J/cm^2^ in a CL-1000 Ultraviolet Crosslinker (UVP), sodium pyruvate was added to a final concentration of 5 mM to prevent damage from free radicals induced by ultraviolet radiation (62).

For NPC infections, confluent cell monolayers on fibronectin-coated dishes were dissociated using Accutase (Millipore), 3×10^6^ cells were reseeded in poly-D-lysine-coated 100-mm dishes or uncoated dishes with poly-D-lysine-coated coverslips, and cells were allowed to attach overnight. Cells were exposed to HCMV at the indicated multiplicities of infection (MOIs). For the evaluation of IE1-directed shRNAs, NPCs or HEL cells were infected with HCMV at an MOI of 1. To overcome the delay of infection process of TNdlIE1, MOI of 10 was used. After incubation for 3 h to allow for virus adsorption, the inoculum was removed and cells were refed with fresh medium. Cells were collected and analyzed at the indicated times post infection. To study infection in the absence of *de novo* protein synthesis, NPCs were pretreated with 10 μg/ml cycloheximide (CHX, Sigma) for 1 h prior to infection, infected with HCMV in the presence of CHX (10 μg/ml), and collected at 16 hpi for RNA and protein analysis.

### Lentivirus preparation and transduction

Stocks of replication-defective lentiviruses were prepared as described previously (57, 63). Briefly, 1.5×10^6^ HEK 293T cells were seeded in a 100-mm dish. On the next day, calcium phosphate precipitation was applied to cotransfect the cells with packaging plasmids pML-A8.9 (12 μg) and pVSV-G (8 μg) (System Biosciences) along with 15 μg of one of the following expression plasmids: pCDH-puro-IE1, pLKO.1-scramble, pLKO.1-shRNA-IE1-1, pLKO.1 -shRNA-IE1 −2, pLKO.1-shRNA-IE1-3, Tet-pLKO-puro-shDsRed, Tet-pLKO-puro-shLuci, Tet-pLKO-puro-shSTAT3-1, or Tet-pLKO-puro-shSTAT3-2. Following a medium change, the lentivirus containing supernatants were collected 72 h after transfection and stored at −80°C.

The lentivirus stocks were used to transduce HEL cells, HELf cells, or NPCs. To this end, 5×10^6^ NPCs were seeded in fibronectin-coated dishes and infected with equal volumes (2 ml) of lentivirus stock on the following day. Lentivirus stock was added to the NPCs again on the next day. The inoculum was replaced with fresh culture medium 3 h after each transduction. Likewise, 5×10^5^ HEL cells were infected with 2 ml lentivirus stock on the day following seeding, and the inoculum was replaced with fresh culture medium 5 h after infection. The transduced cells were cultured for 3 days to allow for transgene expression, before they were subjected to HCMV infection and/or RNA or protein analysis. The transduction procedure used to establish HELf-IE1 cells is described above.

### Gene silencing with shRNAs

To determine the IE1-specific silencing efficiency, equal amounts of shRNA expressing lentiviruses (sh-1, sh-2, sh-3, and scr) were used to transduce HEL cells in parallel. The resulting HEL cells were cultured for 2 days to allow for shRNA expression, and this was followed by 2 days serum starvation with serum free medium to synchronize the cells. The synchronized cells were reseeded in 60-mm dishes (1×10^6^ cells/dish), allowed to attach, and infected with HCMV (MOI = 1). The infected cells were collected at 24 and 48 hpi, and the levels of IE1 were determined by protein analysis. After selection of the most efficient shRNA (sh-2), NPCs were transduced with sh-2 and scr lentiviruses in parallel. Transduced NPCs were cultured for 48 h to allow for shRNA expression, reseeded in poly-D-lysine-coated dishes (3×10^6^ cells/dish), and infected with HCMV (MOI = 1) on the following day. Cells were collected at the indicated times post infection and subjected to protein and RNA analysis.

NPCs transduced with shRNA expressing lentiviruses Tet-pLKO-puro-shDsRed, Tet-pLKO-puro-shLuci, Tet-pLKO-puro-shSTAT3-1, or Tet-pLKO-puro-shSTAT3-2 (described above) were treated with 1 μg/ml doxycycline hyclate (Dox, Aladdin) for 48 h, with a medium change after 24 h, to induce shRNA expression. Cells were collected and analyzed at the indicated times.

### Quantitative reverse transcriptase PCR (qRT-PCR)

Transfected or infected NPCs were collected at the indicated times. Total RNA was isolated using the RNAiso Plus reagent (Takara) followed by RNase-free DNase I treatment (Thermo Fisher Scientific) to remove residual genomic DNA. Equal amounts (500 ng) of DNA-free RNA were reverse transcribed using the PrimeScript RT Reagent Kit (Perfect Real Time; Takara) according to the manufacturer’s instructions. Then, qPCR was performed in a CFX96 Connect Real-Time PCR Detection System (Bio-Rad). Each 20-μl qPCR reaction contained 2 μl RT product, 10 μl 2× iTaq Universal SYBR Green Supermix (Bio-Rad), and 200 nM forward (F) and reverse (R) primers. Primer sequences are shown in Table 1. Amplification was performed by denaturation at 95°C for 5 min, followed by 35 two-step cycles of 95°C for 10 s and 60°C for 30 s. Melting curve analysis was carried out at 95°C for 1 min, 55°C for 1 min, and 55 to 95°C for 10 s. Each reaction was performed in triplicate, and results for the target gene mRNA were normalized to glyceraldehyde 3-phosphate dehydrogenase (GAPDH) using the *2*^ΔΔCT^ method. Three independent experiments were performed, and results are presented as the means ± one standard deviation (SD). Data were statistically evaluated using the Student’s t-test. A *P*-value of ≤0.05 was considered statistically significant.

### Western blotting

At the indicated times, cells were washed in phosphate-buffered saline (PBS), detached with Accutase, collected, counted, and centrifuged. Cell pellets were snap-frozen in liquid nitrogen and stored at −80°C until completion of the time course. Then, cell pellets were lysed in radioimmunoprecipitation assay (RIPA) buffer. Equal amounts of cell lysates were separated by sodium dodecyl sulfate-polyacrylamide gel electrophoresis (SDS-PAGE) and transferred to polyvinylidene fluoride (PVDF) membrane (Millipore). After incubation with the indicated primary and corresponding secondary antibodies, signals were detected using a Chemiluminescence machine, and analyzed by densitometry program (Image J). At least three sets of independent experiments were performed and representative results were shown. HCMV proteins were detected using mouse monoclonal antibodies to IE1 (clone p63-27, IgG2a), IE1/IE2 (CH16), UL44 (IgG1, Virusys), glycoprotein B (gB; clone 27-156, IgG2b), or pp65 (IgG1, Virusys). Cellular proteins were detected using a goat polyclonal antibody to SOX2 (clone L1D6A2, IgG1), a mouse monoclonal antibody to STAT3 (clone 124H6, IgG2a, Cell Signaling Technology), a rabbit polyclonal antibody to pSTAT3 (Y705) (IgG, Cell Signaling Technology), a mouse monoclonal antibody to β-actin (IgG, Santa Cruz Biotechnology), a rabbit polyclonal antibody to GAPDH (IgG, Proteintech), or a rabbit polyclonal antibody to lamin B1 (IgG, Proteintech). Secondary antibodies used were horseradish peroxidase-conjugated sheep anti-mouse IgG (Amersham Bioscience), donkey anti-rabbit IgG (Amersham Bioscience), or donkey anti-goat IgG (Proteintech).

### Immunofluorescence assay

Viral and cellular proteins in cells grown on coverslips were detected by indirect immunofluorescence analysis as described previously (64). Briefly, NPCs were seeded on poly-D-lysine-coated coverslips in uncoated dishes and mock- or HCMV-infected (MOI = 3). Coverslips were collected at the indicated times post infection. UL44, SOX2 or STAT3 were stained with the respective primary antibodies: mouse monoclonal anti-UL44 (IgG1, Virusys), goat polyclonal anti-SOX2 (Santa Cruz Biotechnology), or mouse monoclonal anti-STAT3 (clone 124H6, IgG2a, Cell Signaling Technology). The secondary antibodies included fluorescein-isothiocyanate (FITC)-conjugated donkey anti-goat IgG (Jackson Immunoresearch), Alexa Fluor 488-conjugated goat anti-mouse IgG1 (Molecular Probes), and Alexa Fluor 488-conjugated goat anti-mouse IgG2a (Molecular Probes). Nuclei were counterstained with Hoechst 33342 dye, and coverslips were mounted with anti-fade mounting solution containing paraphenylenediamine (65). Images were obtained using a Nikon Eclipse 80i or Nikon Eclipse Ti-S epifluorescence microscope equipped with a Nikon DS-Ri1 camera and processed using the NIS-Elements F3.0 software.

### Cellular fractionation

Nucleofected or infected NPCs were washed in pre-cooled PBS and collected by scraping. Cytosolic and nuclear fractions were prepared using the Nuclear-Cytosol Extraction Kit (Applygen Technologies) following the manufacturer’s instructions. Briefly, cell pellets were lysed in pre-cooled Cytosol Extraction Buffer A (CEB-A), vortexed, reacted with Cytosol Extraction Buffer B (CEB-B), vortexed, and centrifuged. The resulting supernatant contained the cytosolic fraction. The precipitate was washed with CEB-A, lysed in Nuclear Extraction Buffer (NEB), vortexed, and centrifuged. The resulting supernatant contained the nuclear fraction.

## ACKNOWLEDGMENTS

We appreciate the critical reading from Tom Shenk. We thank the colleagues mentioned in Materials and Methods for providing important reagents. MHL was supported by the Ministry of Science and Technology of China (National Program on Key Basic Research Project 2015CB755600), the National Natural Science Foundation of China (81620108021, 31170155, and 81427801), the Sino-Africa Joint Research Centre (SAJC201605) and a seed grant from the University of Idaho (YDP-764). MN and CP were supported by the Wellcome Trust Institutional Strategic Support Fund, MN was supported by the Medical Research Council (MR/P022146/1) and Tenovus Scotland (T15/38), and CP was supported by the Deutsche Forschungsgemeinschaft (PA 815/2-1).

